# Experimental Evaluation of Methods for Real-Time EEG Phase-Specific Transcranial Magnetic Stimulation

**DOI:** 10.1101/860874

**Authors:** Sina Shirinpour, Ivan Alekseichuk, Kathleen Mantell, Alexander Opitz

**Affiliations:** Department of Biomedical Engineering, University of Minnesota, Minneapolis, MN, USA

**Keywords:** Transcranial magnetic stimulation, Electroencephalography, Closed-loop brain stimulation

## Abstract

Brain oscillations reflect system-level neural dynamics and capture the current brain state. These brain rhythms can be measured noninvasively in humans with electroencephalography (EEG). Up and down states of brain oscillations capture local changes in neuronal excitability. This makes them a promising target for non-invasive brain stimulation methods such as Transcranial Magnetic Stimulation (TMS). Real-time EEG-TMS systems record ongoing brain signals, process the data, and deliver TMS stimuli at a specific brain state. Despite their promise to increase the temporal specificity of stimulation, best practices and technical solutions are still under development. Here, we implement and compare state-of-the-art methods (Fourier based, Autoregressive Prediction) for real-time EEG-TMS and evaluate their performance both *in silico* and experimentally. We further propose a new robust algorithm for delivering real-time EEG phase-specific stimulation based on short prerecorded EEG training data (Educated Temporal Prediction). We found that Educated Temporal Prediction performs at the same level or better than Fourier-based or Autoregressive methods both *in silico* and *in vivo*, while being computationally more efficient. Further, we document a dependency of EEG signal-to-noise ratio (SNR) on algorithm accuracy across all algorithms. In conclusion, our results can give important insights for real-time TMS-EEG technical development as well as experimental design.

## 1. Introduction

Large-scale brain activity undergoes ongoing cyclical changes resulting from neural activity interaction of cellular and circuit properties (Buzsáki and Draguhn, 2004). These neural oscillations represent rhythmic alternations between high and low excitability brain states (Schroeder and Lakatos, 2009). Electroencephalography (EEG) enables us to noninvasively record these brain oscillations, which are capturing the synchronous activity of neural ensembles (Buzsáki et al., 2012). The instantaneous phase of brain oscillations is an important feature of neural processing (Alekseichuk et al., 2016; Jacobs et al., 2007; Maris et al., 2016; Sauseng and Klimesch, 2008; Thut et al., 2012). Thus, it can serve as an indicator of brain excitability to inform the timing of brain stimulation delivery such as Transcranial Magnetic Stimulation (TMS).

TMS is a non-invasive brain stimulation method that induces strong short-lasting electric fields in the brain (Barker et al., 1985). A TMS pulse is generated by passing a high-intensity electrical current through an inductive coil to create a magnetic field that penetrates the skull and induces a secondary electric field in the brain, which modulates neural activity (Hallett, 2007; Opitz et al., 2011). TMS is being increasingly explored as a tool to modulate brain activity for the treatment of neurological and psychiatric disorders (Lefaucheur et al., 2014). The combination of TMS with EEG can improve stimulation protocols by tailoring them to the individual’s ongoing brain state (Bergmann et al., 2016; Thut et al., 2017).

Numerous studies have assessed the relationship between ongoing EEG oscillations and physiological responses to TMS. For instance, Romei et al., 2008 reported that for the same TMS stimuli, phosphene induction is increased for lower prestimulus alpha power in posterior brain regions. Dugué et al., 2011 found a higher probability of TMS-induced phosphenes during the peaks of occipital alpha oscillations. Reports on whether EEG power and/or phase are relevant markers of corticospinal excitability for TMS motor evoked potentials (MEPs) are mixed in the literature e.g. (Berger et al., 2014; Hussain et al., 2019; Keil et al., 2013; Schulz et al., 2014). However, the inferences to be drawn from these studies are limited, as they employed a *post hoc* analysis of EEG data, thus, not allowing a direct causal test of current EEG brain states on TMS responses.

Real-time EEG-TMS methods that deliver stimulation based on the present brain state have the potential to overcome the limitations of previous EEG-TMS studies. Further, they can potentially deliver more efficient brain stimulation and improve research and clinical outcomes (Karabanov et al., 2016; Polanía et al., 2018; Zrenner et al., 2016). Recent implementations of such real-time EEG-TMS systems delivered TMS based on the phase and power of the EEG mu-rhythm to study MEP corticomotor excitability (Madsen et al., 2019; Zrenner et al., 2018). Further, alpha-synchronized TMS over the dorsolateral prefrontal cortex (DLPFC) has been investigated for depression treatment (Zrenner et al., 2019).

While real-time approaches are a promising technology, they come with a set of technical challenges. In particular, it is difficult to extract the instantaneous phase of the recorded EEG signals accurately in real-time. One main problem of any real-time processing system compared to offline methods is the trade-off between speed and accuracy. Since the data needs to be processed within a limited time window, algorithms have to be computationally fast, even at the cost of accuracy. Another problem is that during real-time computations, we are limited by causality in signal processing, which means that future signal samples are not available for processing such as filtering. In signal processing, band-pass filters extract the frequency bands of interest (e.g. alpha, beta). These filters invariably introduce artifacts around the edges of the available time-series. Since only data recorded up until the current time point are available, filtering inevitably distorts the data and its phase at the current moment. Thus, it is necessary to predict the signal and phase based on the currently available undistorted data from the near past, which can also compensate for any hardware delays.

Multiple methods have been proposed for EEG forward prediction. Since continuous EEG signals to a certain degree are nonsinusoidal and nonstationary (Cole and Voytek, 2017; Mäkinen et al., 2005), this is a challenging problem. Two main methods have been suggested for real-time phase-dependent EEG-TMS: 1) Forward prediction in the time domain. Here, the algorithm uses temporal patterns in a short time window (e.g. the last 0.5 seconds of recorded data) in a frequency band of interest to predict the signal waveform for future time periods. One particular implementation is the autoregression model based prediction (Chen et al., 2013). 2) Forward prediction in the frequency domain. Here, the signal is projected into the Fourier domain to capture the main sinusoidal (Mansouri et al., 2017) or wavelet component (Madsen et al., 2019) to be used for phase extrapolation.

While these methods have been successfully implemented for real-time phase prediction, they have certain drawbacks. Commonly used autoregressive (AR) predictions in the temporal domain need careful parameter optimizations, have high computational demand, and need high signal power in the frequency band of interest which can lead to the exclusion of participants (Zrenner et al., 2018). Frequency-domain algorithms, like in Mansouri et al., 2017 are computationally effective and relatively parameter-free, but their accuracy relies on the periodicity and harmonicity of the brain signal.

To overcome existing accuracy-complexity trade-offs, we propose a third, conceptually independent method that is parameter-free, fast, and provides a similar or higher accuracy compared to the existing methods. This algorithm, named Educated Temporal Prediction (ETP) leverages a short training session before the real-time application to learn each individual’s statistical features of the target brain oscillation. Unlike the first two methods that only use data in the current time window for forward prediction, ETP utilizes pre-learned features for faster or more accurate prediction. Since this algorithm is trained on more data (∼minutes) than it has access to during real-time processing (∼ 1 second), it can be more robust to compensate for occasional reductions in signal quality. In our current implementation, the ETP algorithm relies on a basic statistical characteristic of phasic data – the central moment of the inter-peak intervals in the time domain. The goal of this study is to compare all algorithms using *in silico* and *in vivo* real-time implementations.

## 2. Material and Methods

### 2.1. Phase Prediction Algorithms

Several different classes of algorithms to detect and predict the EEG phase for TMS closed-loop stimulation have been proposed in the literature. Below we summarize the key principles for (a) Fast Fourier Transform (FFT) based prediction, (b) Autoregressive (AR) forecasting, and (c) newly suggested Educated Temporal Prediction (ETP).

#### Fast Fourier Transform (FFT) Prediction

The key feature of this algorithm is to use the frequency domain of the EEG signal for forward prediction e.g. (Mansouri et al., 2017). The specific implementation in our experiment (Fig. 1A) is as follows: 1) The Laplacian montage for the desired brain region is applied to the signal (Hjorth, 1980) (Fig. 1E). 2) The signal is zero-phase FIR (Finite Impulse Response) filtered in the frequency band of interest (Alpha: 8-13 Hz, Beta: 14-30 Hz) using the Fieldtrip toolbox (Oostenveld et al., 2010). 3) The FFT of the signal is calculated. 4) The dominant oscillation (maximum amplitude in the frequency spectrum) is detected. 5) The phase of the signal at the dominant frequency is calculated from the angular component of the complex FFT signal. 6) A sine wave of the dominant oscillation with given frequency and phase as calculated in the previous steps is used for forward prediction.

**Fig. 1.**
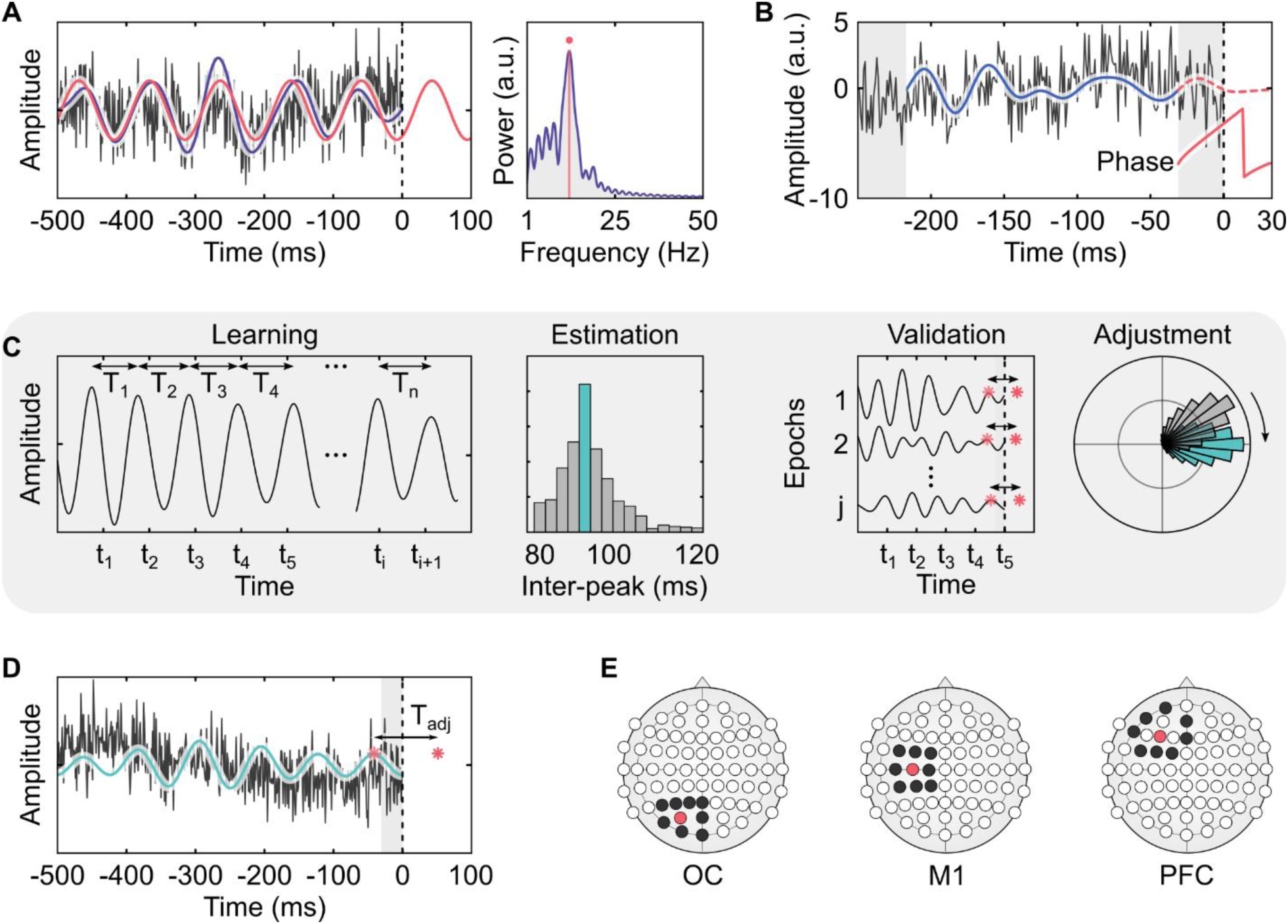
Algorithm implementation. **(A)** FFT algorithm. Left panel: The raw data (black) is band-pass filtered (purple). Right panel: the filtered data is Fourier transformed (purple), the frequency and phase of the dominant frequency are calculated (red). A sine wave with the calculated frequency and phase is used for forward prediction (red, left panel). **(B)** AR algorithm. The raw data (black) is band-pass filtered (blue). Signal edges on both sides are removed (gray). The autoregressive model is estimated and the signal is predicted (dotted red). The phase of the predicted signal is calculated using the Hilbert transform (Solid red). **(C)** ETP training. During the learning phase, the first half of the resting state data is band-pass filtered and the distances between peaks are calculated. The median of the peak periods are calculated as the initial interpeak interval *T*. The second half of the resting-state data is used for the validation phase, the data is segmented into smaller epochs and the peaks are forward projected using *T*. The predicted peaks are compared with the ground truth. Then, *T* is fine-tuned for optimal performance (*T*_*adj*_) **(D)** ETP Real-time application. The raw data (black) is band-pass filtered (green). The edge at the end of the data is removed (gray). The last peak in the time window is projected in the future using *T*_*adj*_. **(E)** Laplacian montages used for the regions of interest. The red electrode is the central electrode. The mean of the surrounding electrodes (black) is removed from the central electrode.

#### AutoRegressive (AR) Prediction

In this method, the signal is predicted in the time domain. A detailed description can be found in (Zrenner et al., 2018; Chen et al., 2013). In our implementation we perform the following steps (Fig. 1B): 1) The Laplacian of the electrodes corresponding to the region of interest is calculated. 2) The signal is zero-phase band-pass filtered in the frequency band of interest using an FIR filter. 3) The edges of the signal are trimmed to remove edge artifacts due to filtering. 4) The remaining signal is used to calculate the coefficients for the autoregressive model. 5) Using the AR coefficients, the signal is iteratively forward predicted. 6) The instantaneous phase of the predicted signal is calculated using the Hilbert transform.

#### Educated Temporal Prediction (ETP)

In this method, we propose to include a short training phase to learn the individual statistical properties of the oscillation of interest. We use a simple and robust method to extract inter-peak intervals and their central moment. Assuming that brain oscillations are quasi-stable over the short measurement periods, one can determine the typical interval between subsequent signal peaks (corresponding to 360° in phase). To predict the time point at which the next target phase, i.e. the peak, will occur, one can add the period between signal peaks to the time of the last peak in the current recorded signal window. Here, we have implemented this algorithm as shown in Fig. 1C-D: 1) Three minutes of resting-state data is recorded. 2) The Laplacian montage for the region of interest is applied. 3) The data is filtered with a zero-phase FIR filter in the band of interest. This signal is used as the ground truth for later steps. 4) Signal peaks in the first 90 seconds of the training data (learning phase) are identified (*T*_*n*_). To ensure that found peaks are meaningful and not a result of noise, a criterion of minimum peak distance (i.e. 62.5 ms for alpha oscillations) was introduced. 5) The median of the periods between those peaks was chosen as the interpeak interval (*T*) to be used for peak prediction. 6) The second 90 seconds of raw resting EEG data are used for the estimation of the accuracy of the phase prediction (validation phase). For this, 250 overlapping windows with a length of 500 ms are selected. 7) The Laplacian montage of the region of interest is applied and the windows are individually filtered in the band of interest using a non-causal sinc filter (brick-wall filter in the Fieldtrip toolbox). 8) The edge (40 ms for alpha) from the end of the window is removed to avoid the edge artifact caused by the filter. 9) The peaks in this window with the criteria explained in step 4 are detected. 10) The timing of the next peak is predicted by adding *T* to the last detected peak. 11) The actual phases of the signal at the predicted peaks are measured from the Hilbert transform of the ground truth. 12) In an ideal case, the actual phase of all the predicted peaks should have no deviation from the target phase. However, due to the presence of noise, the achieved phase values may differ. To minimize the potential error in the phase detection, the value of *T* is increased or decreased incrementally to achieve a zero-mean deviation from the target phase. The adjusted value (*T*_*adj*_) is used for real-time phase estimation. In other words, the bias in phase detection is removed without a change in variance. The calculated phases in this section can also be used as an estimation of the algorithm accuracy in real-time performance. In the real-time segment of the experiment, using *T*_*adj*_, steps 7-10 described above are applied to the current window to predict the next peak (Fig. 1D). If stimulation at any other phase is desired, we can progress the phase of the signal linearly and calculate the time projection needed from the last peak according to equation (1): 

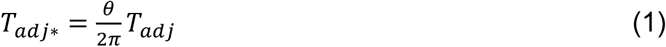

Where *T*_*adj\**_ is the new value of *T* to be added to the last detected peak in the time window, *θ* is the target phase for the stimulation.

### 2.2. *In Silico* Validation Based on Prerecorded Resting-State EEG

#### Algorithm Development and Parameter Optimization

We used a separate prerecorded resting-state EEG dataset for the development and initial optimization of the algorithms. For this, data from 25 individuals (13 female) with an average age of 19 years were taken from the Child Mind Institute healthy brain network dataset (Alexander et al., 2017). Phase detection was simulated on segments of resting EEG data which acted as surrogates of the real-time data. In the development process, we optimized parameters such as filter type, filter order, number of samples for edge removal, peak detection. Priority was given to algorithm performance (i.e. phase accuracy) as long as the process was fast enough to run in real-time. In case the performance of two procedures was equivalent, the faster one was chosen.

#### Algorithm validation

After development, we used another dataset to estimate the accuracy and speed of each algorithm to ensure the generalizability of the results. This dataset included resting-state EEG from 13 healthy adults sampled at 1 kHz with a high number of electrodes (Sockeel et al., 2016). Each individual’s data was epoched with a 500 ms window length and a 50% overlap. We used the three algorithms, FFT, AR and ETP, to predict the phase over three brain regions: left prefrontal cortex (PFC), left motor cortex (M1) and left occipital cortex (OC). These predicted values were compared to the ground truth (continuous EEG data with a two-pass high-order FIR filter and Hilbert transformed to extract phase) to measure the performance. The computation time needed for each algorithm to process the data was also recorded for each epoch.

### 2.3. *In Vivo* Real-Time Experiments

#### System Validation

In order to technically validate the closed-loop system, including hardware and software, we created a test scenario using a dummy head model made of a plastic frame and soft fabric soaked with the saline solution and a known electrical input signal (alternating current at 10 Hz) generated with the XCSITE 100 amplifier (Pulvinar Neuro, Chapel Hill, North Carolina, US). Since the signals recorded by all EEG electrodes are perfect oscillations (within technical limits) with a high signal-to-noise ratio (SNR), we expected to detect the phase correctly using all algorithms (refer to Fig. S1 for results).

#### Human Participants Testing

The study protocol was approved by the Institutional Review Board of the University of Minnesota. All volunteers gave written informed consent prior to participation. Eight healthy participants (three female, average age = 27) without a history of neurological disorders took part in the main experiment, where we targeted the alpha oscillations. In one participant, due to a high level of noise during the measurement, the recordings of the motor region for all the methods were removed. In the main experiment, we targeted the alpha band (8 - 13 Hz). To test whether the algorithms can perform well in a different frequency band, four participants (two females) were called back to test the beta band condition. For this test, we changed the band-pass filter frequency to (14-30 Hz), while keeping all the other parameters the same.

#### Experimental Protocol

The volunteers were asked to sit in a comfortable chair in a relaxed position throughout the experiment. The participants were instructed to maintain a resting state with eyes open during the recordings. The experimental paradigm is illustrated in Fig. 2. Five minutes of resting-state were recorded at the beginning and at the end of the session. We tested the three algorithms to predict the EEG alpha phase over three brain regions (left PFC, left M1, left OC) in a total of nine blocks, of five minutes each. The order of regions and the order of algorithms within each region was randomized. For the ETP method, there was a separate training block in which three minutes of resting-state data were recorded immediately prior to the real-time block.

**Fig. 2.**
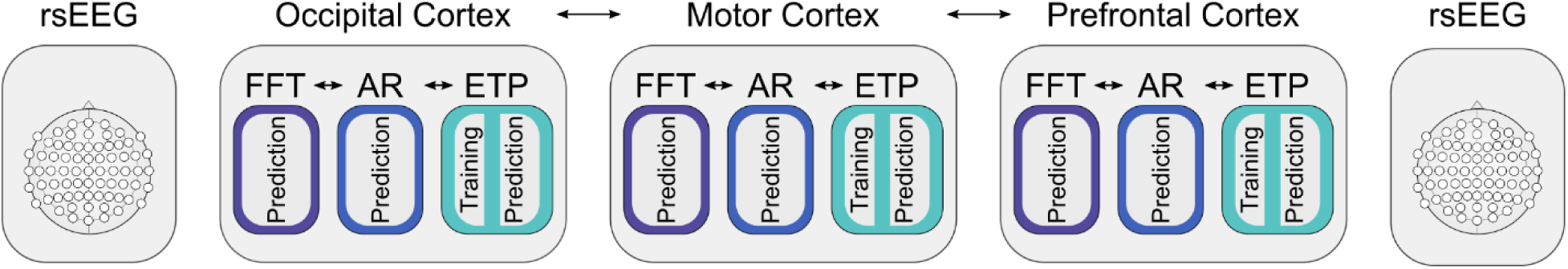
Experimental Protocol. Resting-state EEG is recorded at the beginning and end of the experiments. For each three brain regions that are randomized in order, three algorithms are tested in real-time. Prior to the real-time ETP block, resting-state is recorded and the algorithm is trained.

#### Electrophysiological Recordings

Scalp EEG was recorded using a 64-channel EEG amplifier (actiCHamp, Brain Products GmbH, Germany) with active Ag/AgCl electrodes (actiCAP slim, Brain Products GmbH, Germany). The data were recorded at a sampling rate of 10 kHz with 24 bits resolution. The impedance between the scalp and each electrode was kept below 20 kΩ.

Since we are interested in recording the signal coming from a local brain region, we used the Laplacian montage that subtracts the mean values of the surrounding electrodes from the electrode of interest. This allows for common-mode rejection of signals coming from sources outside of the region of interest. Fig. 1E illustrates the exact montages that were used for this study.

#### Real-time Digital Signal Processing

EEG data from the amplifier was streamed to the processing computer (Microsoft Windows 10, 4 cores 3.60 GHz CPU, 16 GB RAM) using the Lab Streaming Layer (LSL, https://github.com/sccn/labstreaminglayer) software in real-time. Custom scripts (https://github.umn.edu/OpitzLab/CL-phase) were used in MATLAB R2017b to receive and process the EEG data and send the triggers to the TMS machine (Magstim Rapid^2^, UK). The last 500 ms of the streamed data was selected and fed to the algorithms to perform the phase estimation and prediction. The 500 ms window was updated upon receiving each new sample. In order to perform accurate real-time analysis, it is essential to process the current window of data before the next sample is received. This ensures that the system runs smoothly and sustains real-time performance during the whole session. Thus, the streaming data were downsampled before performing the analysis on each window. Since the ETP and FFT methods are computationally fast, we downsampled the EEG data to 1 kHz. For the AR algorithm, due to high computational demand and consistency with previous work (Zrenner et al., 2018), the data were downsampled to 500 Hz.

#### Real-Time Stimulation Triggers

During the real-time phase estimation, a TTL (Transistor-transistor logic) pulse was sent from the parallel port of the computer to trigger the TMS machine. Due to an overall processing delay (∼1-2 ms) and a lag of the TMS machine to deliver the magnetic pulse from the time it receives the trigger, there is a total trigger delay one needs to account for. Since this delay is stable over time, it was experimentally measured during the system validation and adjusted in the code. Note that the total delay is system-specific and should be measured for each system separately. In our experiments, triggers were sent when the estimated phase approached the proximity of the desired phase adjusted with the technical delay. Since TMS pulses are very strong compared to the EEG signals, they cause large artifacts that would distort the phase estimation to determine the ground truth in offline processing (Ilmoniemi and Kičić, 2009; Herring et al., 2015). Therefore, in this study, since we are mainly interested in comparing the accuracy of different methods, we did not apply a TMS pulse and only recorded the time at which the triggers were sent. This way, the EEG signal was not distorted, allowing us to estimate the ground truth phase for algorithm comparison.

### 2.4. Data Analysis and Statistics

#### Algorithm Performance

We quantified the performance of each algorithm calculating its bias, variance, and accuracy. Bias shows the typical difference of the outcome from the target. Here, we used the average difference between the estimated phase and the target phase. Variance quantifies how spread the outcome distribution is. We used the standard deviation to report to quantify the spread. We defined the accuracy in equation (2): 

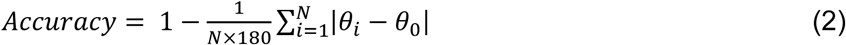

Where *N* is the number of trials for which the phase is estimated, *θ*_*i*_ is the estimated phase (°) for the trial *i*, and *θ*_*0*_ (°) is the desired phase. Accuracy of 1 means that the phase has been estimated precisely all the time. Accuracy of 0.5 is a uniformly random phase estimation. Accuracy of 0 means that all phases have detected as the opposite of the target phase (e.g. troughs instead of peaks).

#### Statistical Comparison Between the Algorithms

To test whether there is a statistical difference between the algorithm performance, we used a general linear mixed-effects model (GLME) with accuracy as the dependent variable, algorithm type as the fixed effect, and brain region as the random effect variables. In case the result of GLME was significant, we used the Wilcoxon signed-rank test to compare each pair of algorithms.

#### Signal-to-Noise Ratio (SNR) Measurements and Accuracy Regression

One key factor for closed-loop performance is the signal-to-noise ratio in EEG recordings. Especially the presence or absence of a prominent alpha rhythm has been used as participant selection criteria in previous studies (Zrenner et al., 2018). We hypothesized that phase estimation accuracy will depend on data quality and alpha power. To test this, we calculated the SNR for each experimental EEG block during post-processing. SNR was measured by calculating the Power Spectral Density (PSD) using Welch’s method with 2s epochs and Hamming windows and then dividing the total power in the alpha band by the total power over all frequencies (1-45 Hz). To quantify this relationship, we calculated the linear regression model between the SNR and phase accuracy of each experimental block for the three algorithms separately.

#### Resting-State Analysis

The ETP algorithm expects that EEG oscillations in the band of interest are quasi-stable over the experiment (in our case, approx. 1 h). To test this, we compared the resting-state data recorded pre-experiment with post-experiment data. Resting-state data were preprocessed to remove noise and artifacts (Delorme and Makeig, 2004; Oostenveld et al., 2010). We calculated the PSD and identified the frequency of maximum PSD for each volunteer in pre and post data. We used the Wilcoxon signed-rank test to evaluate any possible systematic shifts in the alpha peak frequency for all of the three brain sites throughout the experiment.

## 3. Results

### 3.1. *In Silico* Validation Based on Prerecorded Resting-State EEG

First, we evaluated the performance of each algorithm *in silico*. Fig. 3A shows the polar histograms of the difference between the detected phase and the desired phase in the alpha band for FFT, AR, and ETP methods. Fig. S4 also illustrates the polar histograms separately for each brain region. Qualitatively, all algorithms target zero degrees correctly; however, ETP shows the least spread, thus, higher in accuracy, while FFT shows the most spread around the target phase. Additionally, a difference in performance accuracy can be seen between brain regions with occipital cortex showing the highest accuracy followed by M1 and finally PFC (Fig. 3B). Table 1 summarizes the performance metrics (mean, standard deviation, and accuracy) for all algorithms and brain regions. As can be seen, all methods have a negligible bias in phase estimation; however, AR (mean accuracy = 68%) and ETP (71.8%) algorithms manifest lower spreads and therefore higher accuracies compared to the FFT (57.2%) algorithm (by 10.8% and 14.6% higher accuracy, respectively). ETP performs slightly better than AR by 3.8% in terms of accuracy. The general linear mixed effect model confirms significant differences between the algorithms accuracy (F(2,114) = 235, p < 1e-40). Non-parametric pairwise tests indicate a statistically significant difference between the accuracy of each algorithm pairs (AR vs. ETP: p < 1.e-6, AR vs. FFT: p < 1e-7, ETP vs. FFT: p < 1e-7).

**Table 1.** Summary of the performance metrics and computation time for AR, ETP, and FFT algorithms over the brain regions for the *in silico* dataset in the alpha band.

| Algorithm | Region | Mean (°) | SD (°) | Accuracy | Median Time (ms) |
| --- | --- | --- | --- | --- | --- |
| AR | All | 5.44 | 73.59 | 0.68 | 0.78 |
|  | PF | 5.46 | 79.30 | 0.65 | 0.79 |
|  | M1 | 6.37 | 75.88 | 0.67 | 0.81 |
|  | OC | 4.49 | 65.59 | 0.73 | 0.74 |
| ETP | All | 0.37 | 67.35 | 0.72 | 0.06 |
|  | PF | 0.16 | 73.54 | 0.69 | 0.06 |
|  | M1 | 0.38 | 69.49 | 0.71 | 0.06 |
|  | OC | 0.56 | 59.01 | 0.76 | 0.06 |
| FFT | All | 5.55 | 92.36 | 0.57 | 0.82 |
|  | PF | 4.91 | 98.20 | 0.54 | 0.82 |
|  | M1 | 5.92 | 94.33 | 0.56 | 0.82 |
|  | OC | 5.83 | 84.57 | 0.62 | 0.81 |

**Fig. 3.**
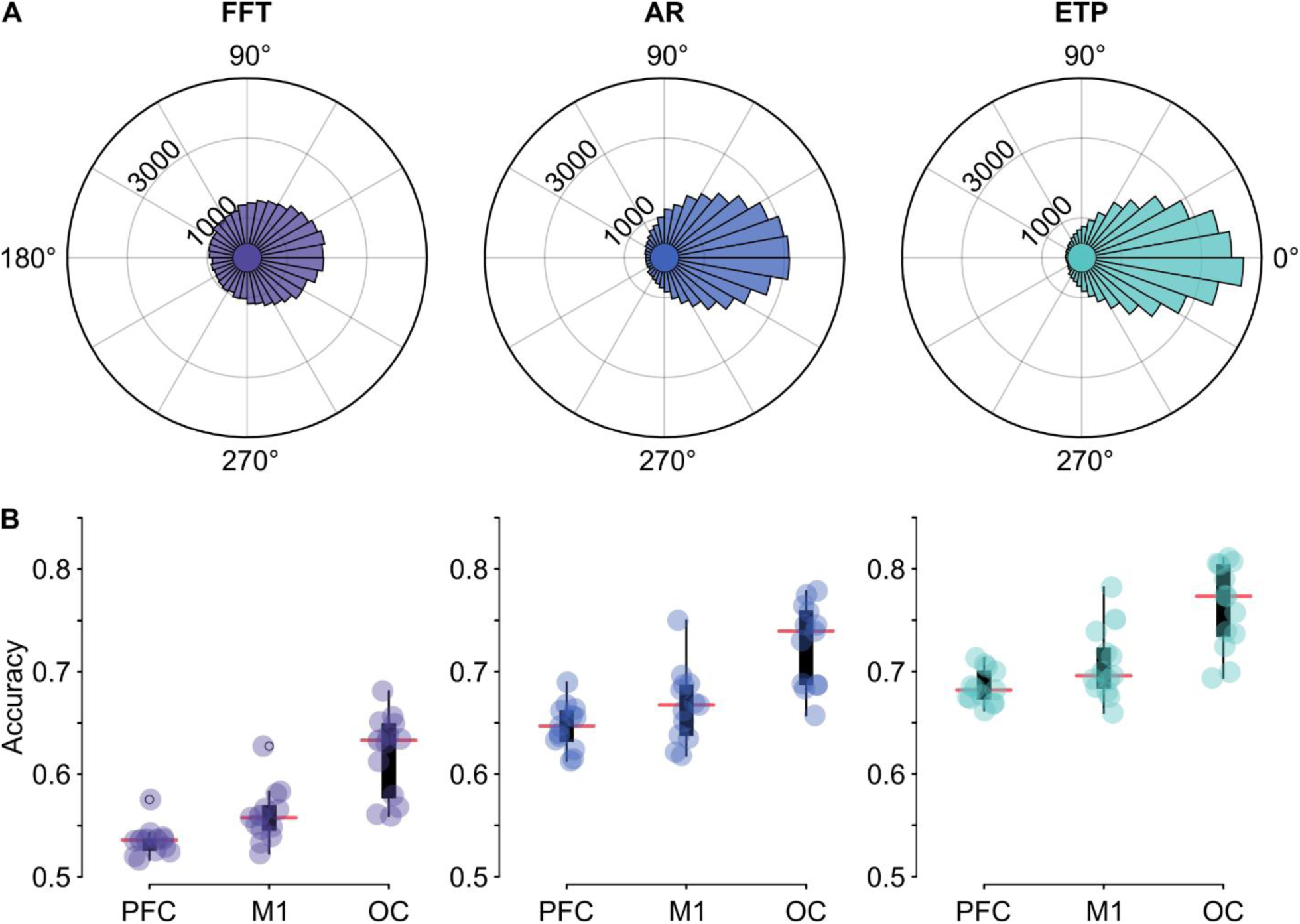
Phase estimation results for the *in silico* dataset in the alpha band. **(A)** Polar histograms of the difference between the estimated phase and ground truth for each algorithm. The phase values are binned into 36 bins. A zero degrees phase would be the ideal outcome since the detected phase matches the desired phase. **(B)** Box plot of the accuracy measurements with the individual datapoints over the three brain sites for each algorithm.

Since computation speed is essential for real-time performance, we also report the median computation times for each method in Table 1. ETP works significantly faster than the two other methods. FFT and AR perform at a similar speed, although the AR method runs at a lower sampling rate which means it is inherently slower. Note that the computation times are hardware-specific and, in order to ensure that the system can run in real-time, the processing time should be kept below the sampling frequency.

To test how well the algorithms generalize, we additionally performed phase estimation in the beta band. Since beta oscillations are faster, less stationary and cover broader frequency band, all algorithms perform worse compared to the alpha band results (Fig. S2 + S5 and Table S1). However, the same pattern between the algorithms still holds. ETP (mean accuracy = 64.7%) shows 3.5% higher accuracy than AR (61.2%), and AR is 5.7% more accurate than FFT (55.5%). The results for computation speeds are comparable to that of the alpha band.

### 3.2. Real-Time Experiments

After comparing the algorithms *in silico* using prerecorded EEG data, we implemented them in the laboratory for true real-time validation. Similar to the *in silico* validation, for the *in vivo* tests, we targeted the peak (zero phase) of the alpha rhythm. The polar histograms are illustrated in Fig. 4A (for separate polar histograms per each brain region see Fig. S6). Fig. 4B breaks down the accuracy of each algorithm over different regions with box plots. The real-time results further validate the findings reported in the *in silico* results, with ETP (mean accuracy = 70.2%) performing marginally better than AR (66.6%), while ETP and AR perform considerably better than FFT (61.1%). Table 2 summarizes the performance metrics of each method in terms of bias, standard deviation, and accuracy for each brain region. As can be seen, ETP shows 3.6% and 9.1% higher accuracy relative to AR and FFT, respectively. Also, AR accuracy is 5.5% higher than FFT. General linear mixed effect model shows significant differences between the algorithms accuracy (F(2,66) = 31, p < 1e-9). Non-parametric pairwise tests confirm that there is a staristaically significant difference between the accuracy of each algorithm pairs data (AR vs. ETP: p < 1.e-3, AR vs. FFT: p < 1e-4, ETP vs. FFT: p < 1e-04).

**Table 2.**
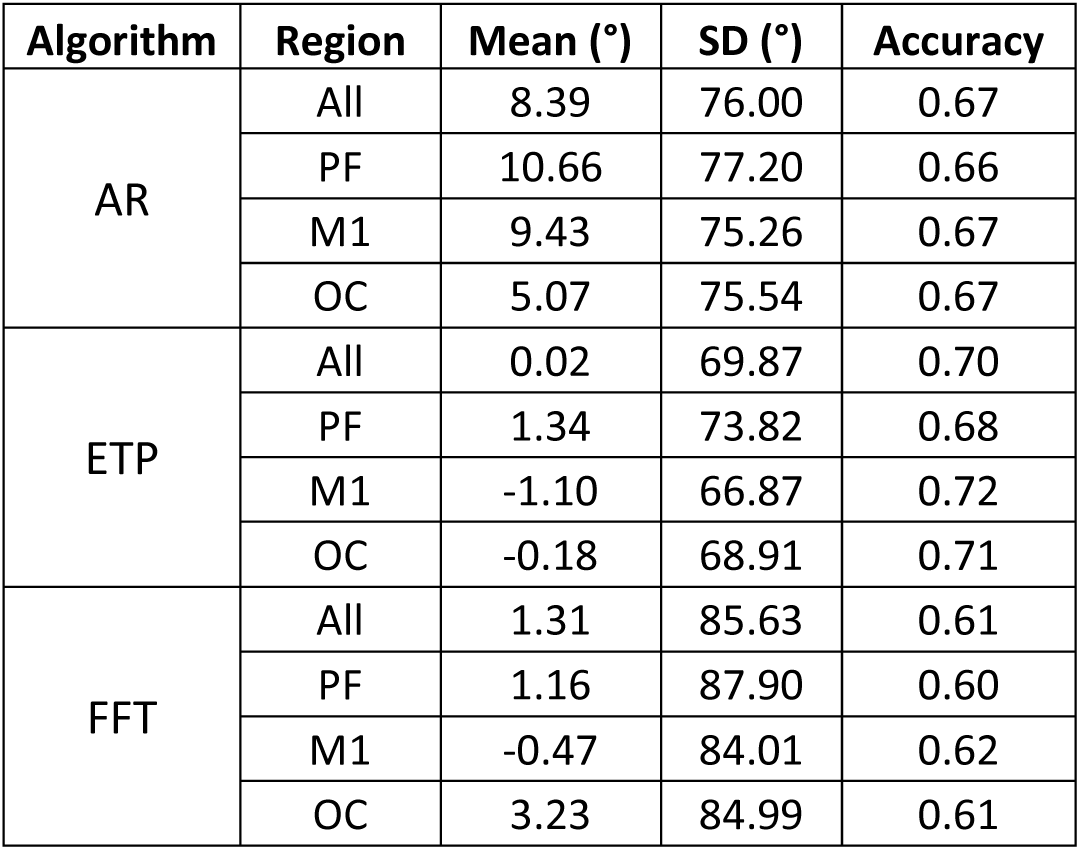
Summary of the performance metrics for AR, ETP, and FFT algorithms over the brain regions for the real-time experiment in the alpha band

| Algorithm | Region | Mean (°) | SD (°) | Accuracy |
| --- | --- | --- | --- | --- |
| AR | All | 8.39 | 76.00 | 0.67 |
|  | PF | 10.66 | 77.20 | 0.66 |
|  | M1 | 9.43 | 75.26 | 0.67 |
|  | OC | 5.07 | 75.54 | 0.67 |
| ETP | All | 0.02 | 69.87 | 0.70 |
|  | PF | 1.34 | 73.82 | 0.68 |
|  | M1 | -1.10 | 66.87 | 0.72 |
|  | OC | -0.18 | 68.91 | 0.71 |
| FFT | All | 1.31 | 85.63 | 0.61 |
|  | PF | 1.16 | 87.90 | 0.60 |
|  | M1 | -0.47 | 84.01 | 0.62 |
|  | OC | 3.23 | 84.99 | 0.61 |

**Fig. 4.**
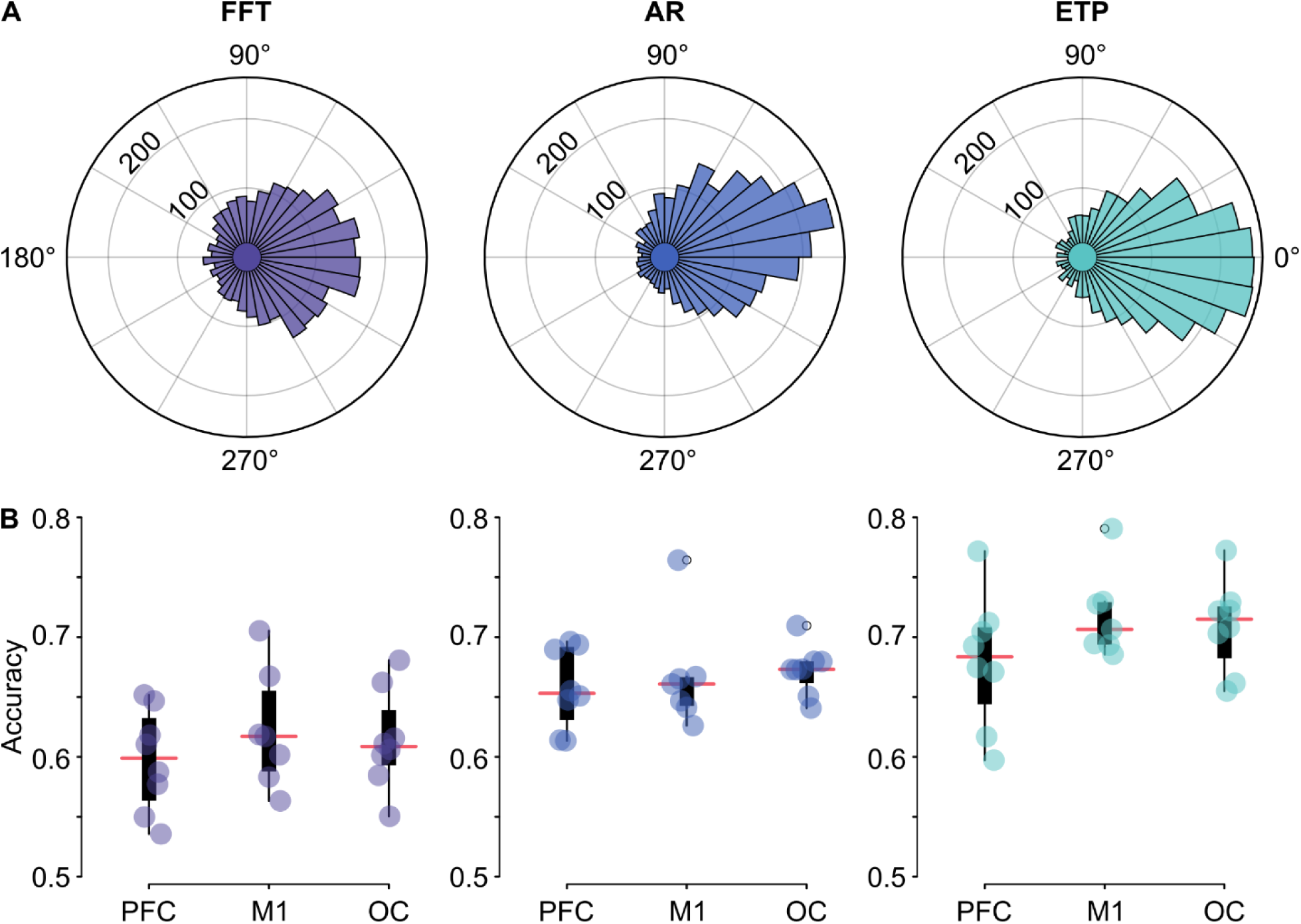
Phase estimation results for the real-time experiment in the alpha band. **(A)** Polar histograms of the difference between the estimated phase and ground truth for each algorithm. **(B)** Box plot of the accuracy measurements with the individual datapoints over the three brain sites for each algorithm.

We also evaluated a real-time phase estimation in the beta band. The polar histograms for beta phase estimations can be found in Fig. S3 and Fig. S7. ETP (mean accuracy = 65.4%) performs considerably better in the beta band as opposed to the other methods with 6.7% and 10% higher accuracy than AR (58.7%) and FFT (55.4%), respectively. These results are expected because the FFT method relies on finding the dominant oscillation to predict the signal and the beta band doesn’t have such stable narrowband oscillaions as the alpha rhythm. For the AR method, optimal parameters such as filter order, edge removal, and autoregressive order are highly dependent on the frequency band of interest. Parameters used here were based on previous research (Zrenner et al., 2018) for the µ-rhythm in the motor cortex and tuned for the frequencies similar to the alpha band. Thus, it is likely that AR will need to be adapted for different frequencies for optimal performance. Our results indicate that ETP, which is a parameter free method, is readily extendable to different EEG rhythms.

#### Signal-to-Noise Ratio (SNR) Measurements and Accuracy Regression

To evaluate the importance of SNR for the phase prediction performance, we applied a linear regression model between the algorithm accuracies and SNR during the measurement. We defined SRN as the ratio of the alpha power to the total power of the signal. Fig. 5 illustrates the regression results for each algorithm. In summary, all methods show a statistically significant increase in phase estimation accuracy for higher SNRs (FFT: R^2^_adj_ = 0.54, F(21) = 26.3, p < 0.01; AR: R^2^_adj_ = 0.41, F(21) = 16.3, p < 0.01; ETP: R^2^_adj_ = 0.44, F(21) = 18.4, p < 0.01). The regression slopes are comparable between the methods (0.31 for FFT, 0.28 for AR, and 0.32 for ETP).

**Fig. 5.**
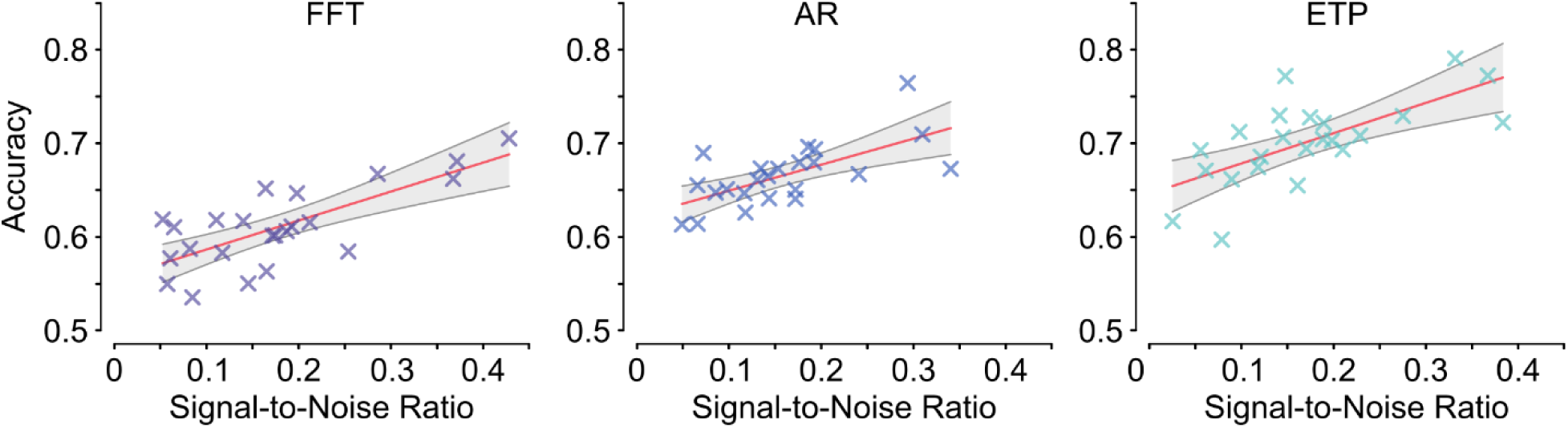
Dependence of phase accuracy during the alpha band stimulation on SNR for each algorithm. The linear regression lines are shown in red and the confidence intervals in grey.

#### Resting-state EEG Analysis

To test whether EEG oscillations are stable throughout the experiment, we compared the frequency of maximum PSD in the alpha band between the pre- and post-experiment resting-state recordings across participants for all three sites using non-parametric paired test. We found no systematic differences (Prefrontal region: p-value = 0.53, Motor: p-value = 0.99, Occipital: p-value = 0.58; all degrees of freedom = 22).

## 4. Discussion

Real-time EEG based TMS applications have high potential to develop more precise stimulation protocols tailored to the individual’s ongoing brain state. However, optimal technical solutions are still being developed and evaluated. We introduced a new education-based method in which the algorithm learns important features from a short prerecorded EEG session. We further compared the accuracy of three conceptually distinct algorithms (temporal prediction, frequency domain prediction, and education-based prediction) to target the peaks of the alpha rhythm over three different brain regions. We compared the performance of these algorithms first *in silico* using prerecorded EEG data and second *in vivo* real-time experiments in human participants. We found the FFT method to perform least accurate among the three studied algorithms, likely due to reducing the signal to a single-frequency sine wave in the observed time window. Since EEG signals are non-stationary and composed of several frequency components, such simplification can impair forward prediction. Our proposed education-based algorithm (ETP) outperformed the two other methods but was otherwise close in accuracy to the AR method. The performance of the ETP method might be due to the robustness of the algorithm since it learns EEG signal patterns beforehand and doesn’t rely only on a short and possibly noisy segment of data that it has access to at the moment of online processing. In addition, we did not exclude any volunteer or data epoch to evaluate performance across a more representative sample. However, as indicated by the found correlation between algorithm accuracy and SNR, higher accuracies could be achieved by excluding participants or time windows with low SNR or oscillatory power.

For real-time applications, computation speed is essential. Hence, we compared the run-time speed of all three algorithms. For a given sampling rate, the AR algorithm is the most computationally demanding. This is due to the heavy calculations needed to compute autoregressive coefficients. On the other hand, because of the simplicity of the ETP method, its computation time is significantly lower than the other two methods. Thus, ETP can run on lower-end hardware and is more accessible to use. Another upside to fast computation time is that it provides more room for future developments in real-time processing. One drawback of using the ETP method is that it needs to be trained on prerecorded data. However, in our case three minutes of resting-EEG data were fully sufficient for training, which can be easily integrated in a real-time experiment. Further, the ETP algorithm is computationally robust and does not need to tune several parameters such as the AR method. For example, the order of the autoregressive model used in the AR algorithm is highly dependent on the sampling rate and frequency of interest on each application, and no trivial method for this optimization is available (Krusienski et al., 2006; McFarland and Wolpaw, 2008). For all three algorithms used, filter type and order can strongly affect the performance and careful consideration is essential for their proper optimization along with any algorithm-specific parameters.

A widespread implementation of real-time TMS-EEG has several potential advantages for future research. For example, brain-state dependent EEG-TMS studies can remove the need for *post hoc* analysis since the brain state of interest can be controlled during the experiments. Furthermore, closed-loop systems using neural biomarkers can be investigated in clinical applications to improve brain stimulation treatments. For instance, repetitive TMS (rTMS) to the left DLPFC is widely used for treating medication-resistant major depression (Avery et al., 2006; O’Reardon et al., 2007). Currently, the treatment protocol does not consider the current brain state. However, it is known that EEG oscillations play an important role in major depression which can be utilized for treatment plans (Fingelkurts and Fingelkurts, 2015; Leuchter et al., 2015). Timing TMS to a high-excitable state might mean that the brain will be more responsive (Zrenner et al., 2019), possibly leading to better treatment outcomes or shorter treatment duration.

Beyond the algorithms evaluated in this study other less common methods to estimate the signal phase exist. (Madsen et al., 2019) used wavelet transforms, which are conceptually similar to the FFT method. Recently, machine learning algorithms have been suggested to estimate the instantaneous phase from unfiltered EEG data (McIntosh and Sajda, 2019). This is similar to the ETP method since it uses prerecorded data to extract features which can inform the real-time application. However, all existing algorithms have room for improvement in accuracy. Due to challenges in implementing real-time phase estimation (such as causality, filtering edge artifact, and system delay), further efforts are needed to improve the performance of such online phase estimation and prediction algorithms. Here, we propose a simple and parameter-free education-based method that only uses easy to access temporal features in the signal (i.e. inter-peak intervals). In the future, more sophisticated algorithms combining temporal, frequency, and spatial features present in resting-state EEG can be developed for more accurate phase prediction in real-time applications. Due to the increasing adoption of brain state-dependent neuromodulation approaches in research and clinical applications, further technical developments can help reach the full potential of this emerging field.

## Acknowledgments

Research presented here was supported by the University of Minnesota’s MnDRIVE Initiative.

## Supplementary Data

**Table S1.** Summary of the performance metrics and speed for AR, ETP, and FFT algorithms over the three brain regions for the *in silico* dataset in the beta band.

| Algorithm | Region | Mean (°) | SD (°) | Accuracy | Median Time (ms) |
| --- | --- | --- | --- | --- | --- |
| AR | All | 0.04 | 85.75 | 0.61 | 0.69 |
|  | PFC | 0.78 | 85.29 | 0.62 | 0.69 |
|  | M1 | 0.57 | 84.65 | 0.62 | 0.69 |
|  | OC | -1.23 | 87.30 | 0.60 | 0.69 |
| ETP | All | 0.81 | 79.84 | 0.65 | 0.08 |
|  | PFC | 1.26 | 79.98 | 0.65 | 0.09 |
|  | M1 | 0.27 | 79.31 | 0.65 | 0.08 |
|  | OC | 0.92 | 80.24 | 0.64 | 0.08 |
| FFT | All | 2.17 | 95.06 | 0.56 | 0.83 |
|  | PFC | 3.51 | 95.64 | 0.55 | 0.83 |
|  | M1 | 2.82 | 94.02 | 0.56 | 0.83 |
|  | OC | 0.16 | 95.53 | 0.55 | 0.83 |

**Table S2.** Summary of the performance metrics for AR, ETP, and FFT algorithms over the three brain regions for the real-time experiment in the beta band.

| Algorithm | Region | Mean (°) | SD (°) | Accuracy |
| --- | --- | --- | --- | --- |
| AR | All | -0.17 | 89.36 | 0.59 |
|  | PFC | -4.18 | 87.82 | 0.60 |
|  | M1 | -3.93 | 90.34 | 0.58 |
|  | OC | 7.59 | 89.92 | 0.59 |
| ETP | All | -2.17 | 78.51 | 0.65 |
|  | PFC | -6.03 | 77.22 | 0.67 |
|  | M1 | -2.80 | 80.25 | 0.64 |
|  | OC | 2.33 | 78.05 | 0.66 |
| FFT | All | -5.09 | 93.70 | 0.55 |
|  | PFC | -3.22 | 91.47 | 0.58 |
|  | M1 | -9.46 | 95.17 | 0.54 |
|  | OC | -2.58 | 94.46 | 0.55 |

**Figure S1.**
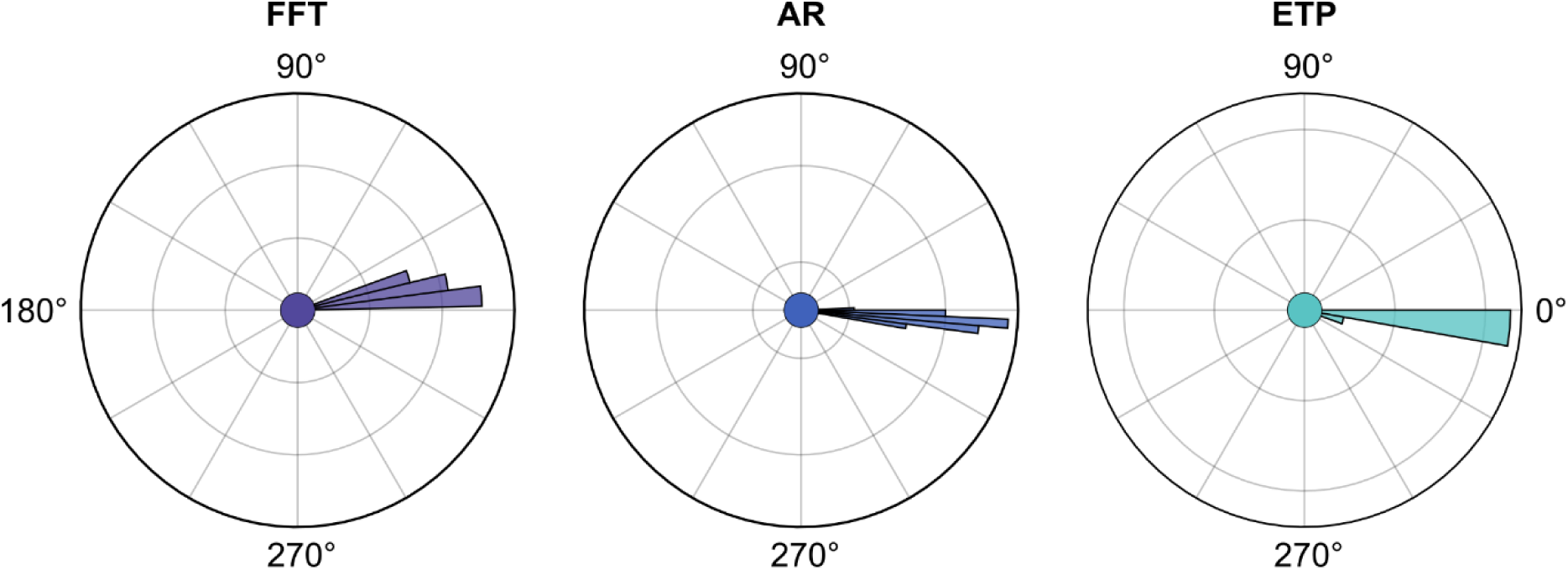
Phase estimation results of the dummy head experiment for the system validation.

**Figure S2.**
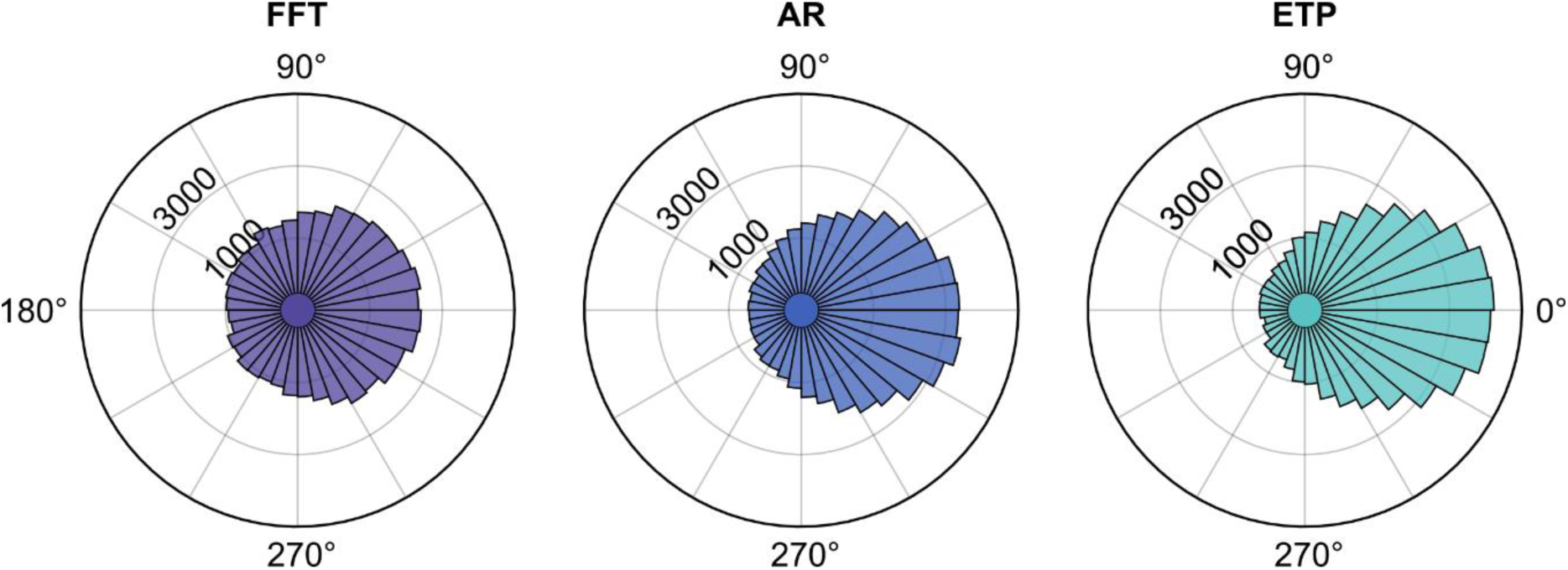
Phase estimation results for the *in silico* dataset in the beta band.

**Figure S3.**
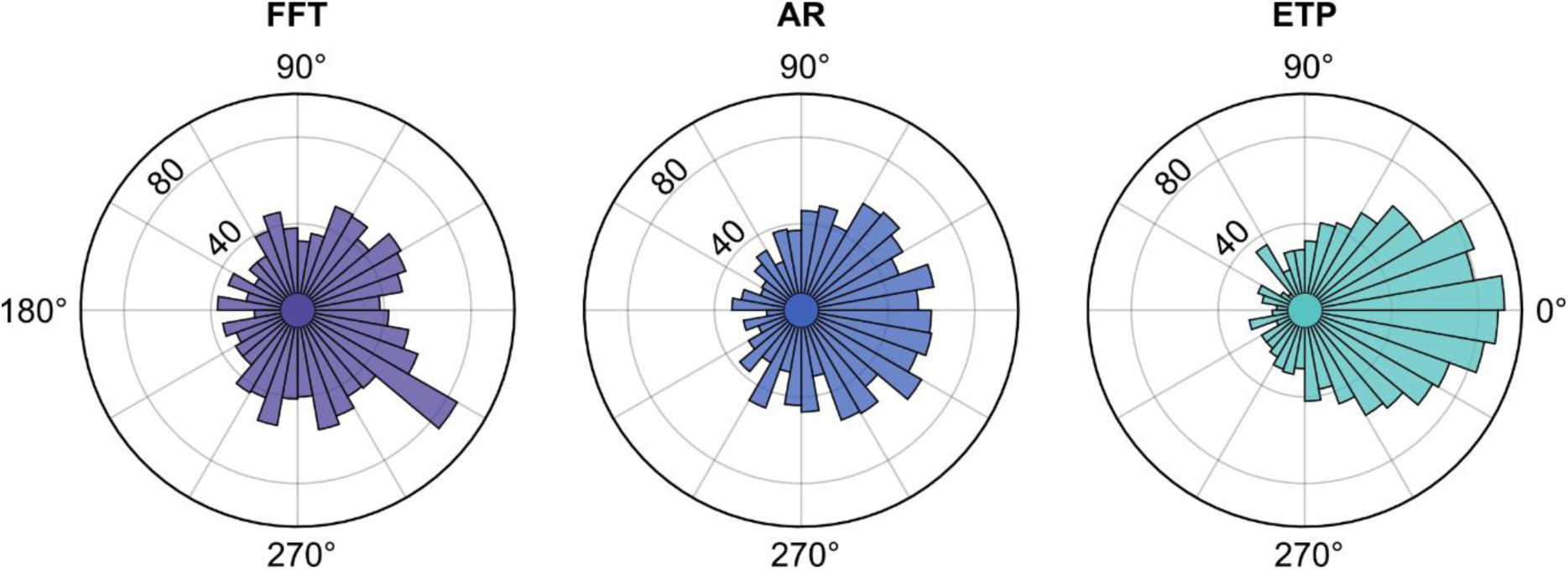
Phase estimation results for the real-time experiment in the beta band.

**Figure S4.**
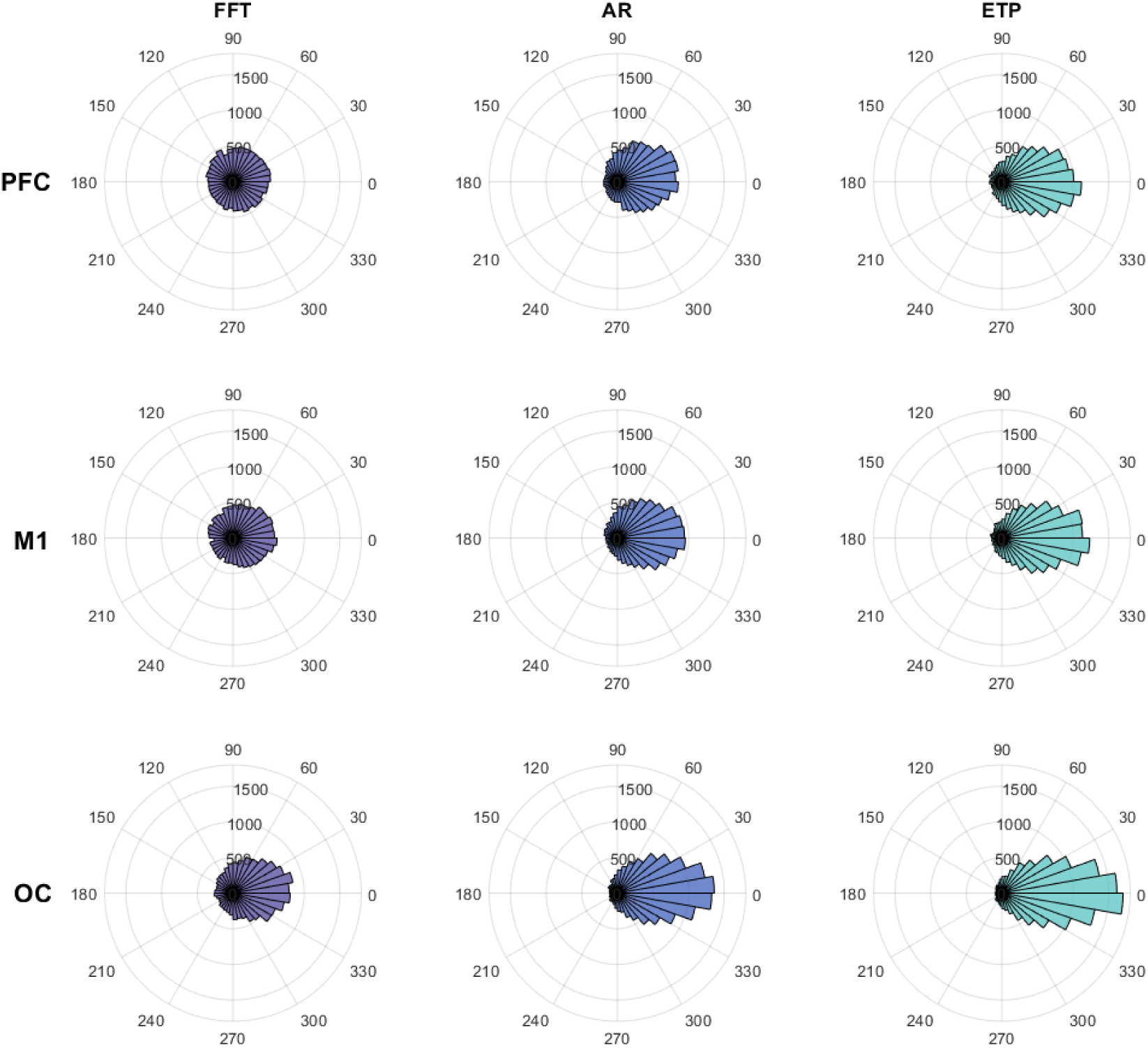
Phase estimation results for the *in silico* dataset for each algorithm and brain region in the alpha band.

**Figure S5.**
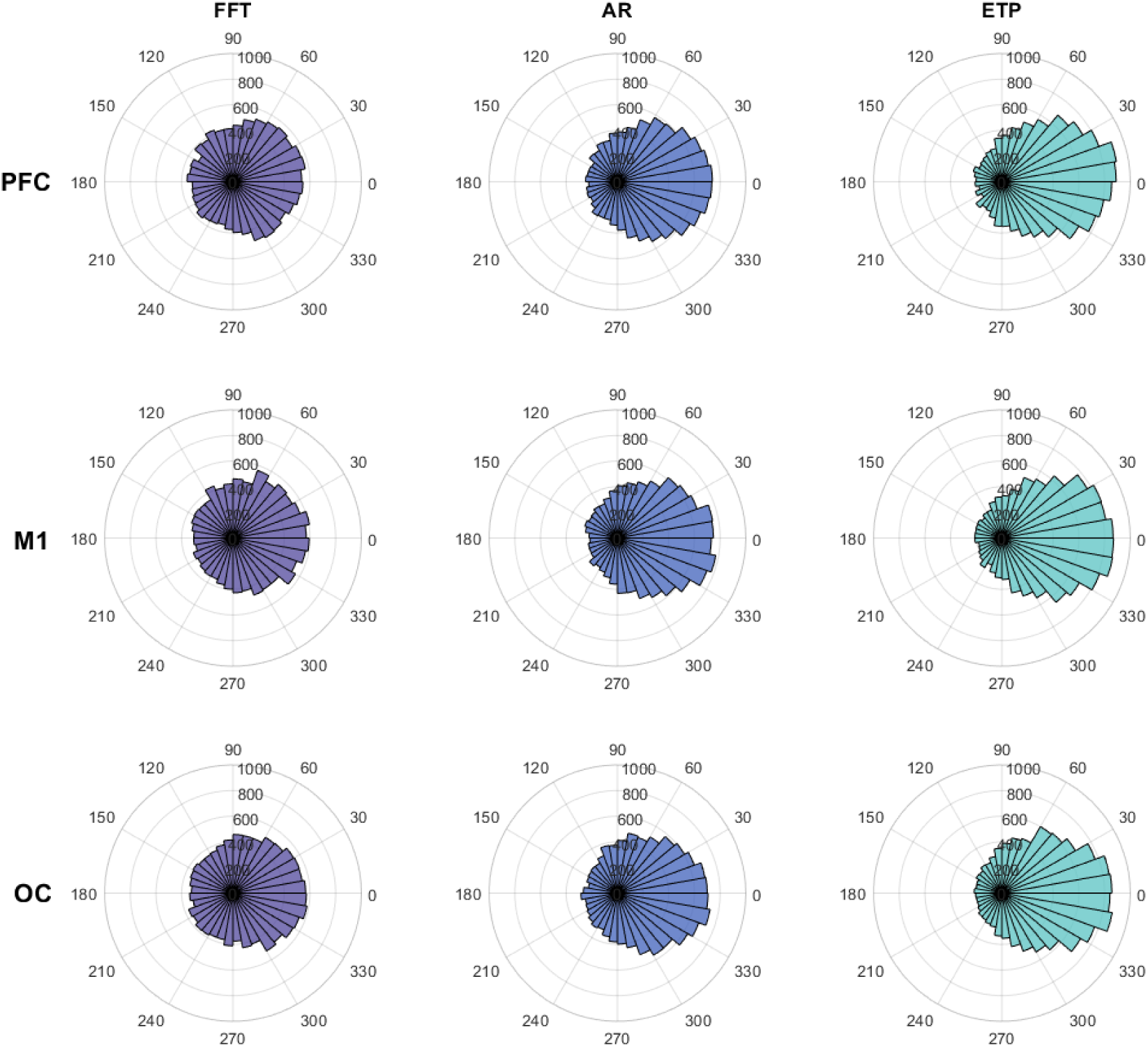
Phase estimation results for the *in silico* dataset for each algorithm and brain region in the beta band.

**Figure S6.**
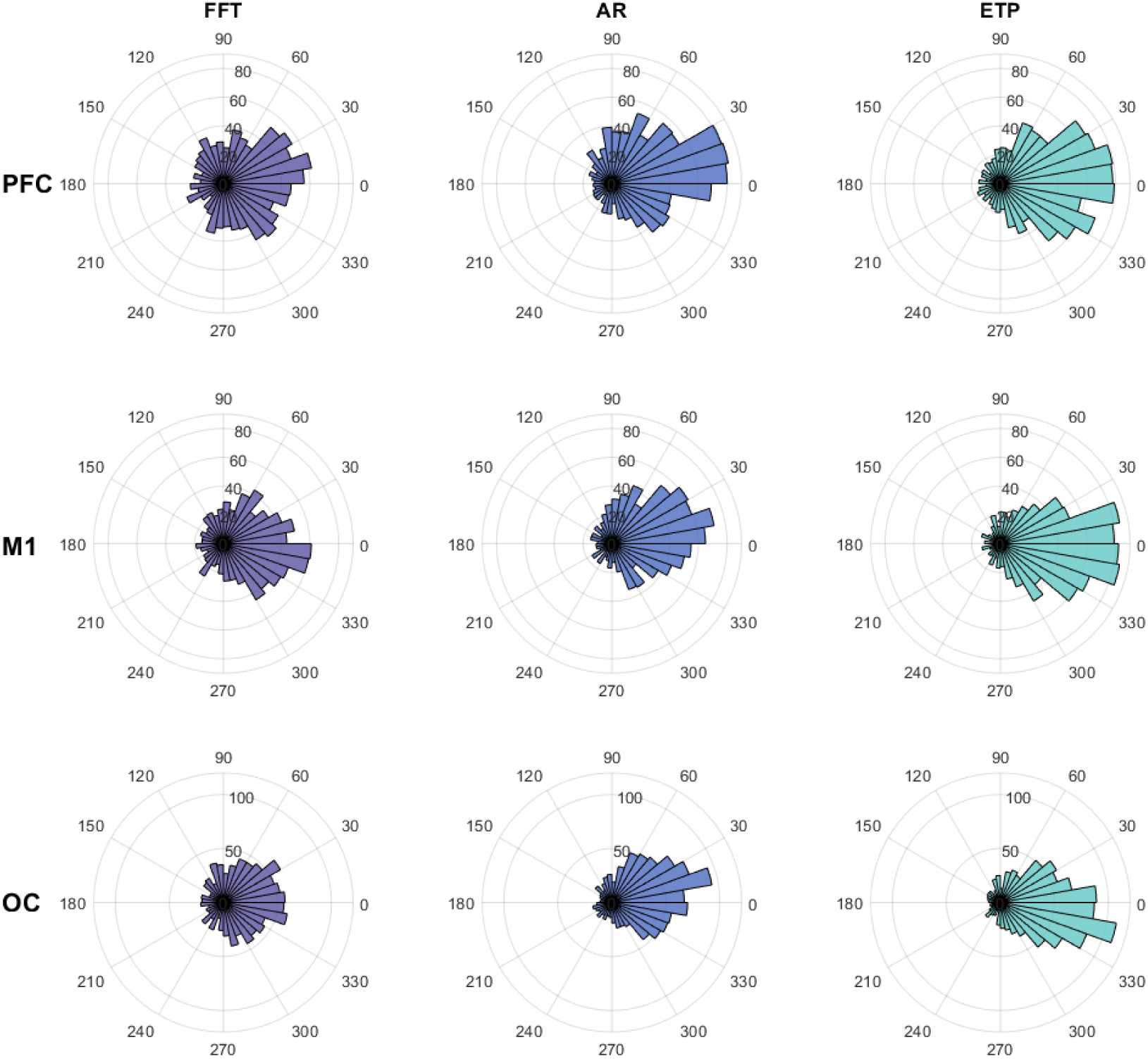
Phase estimation results for the real-time experiment for each algorithm and brain region in the alpha band.

**Figure S7.**
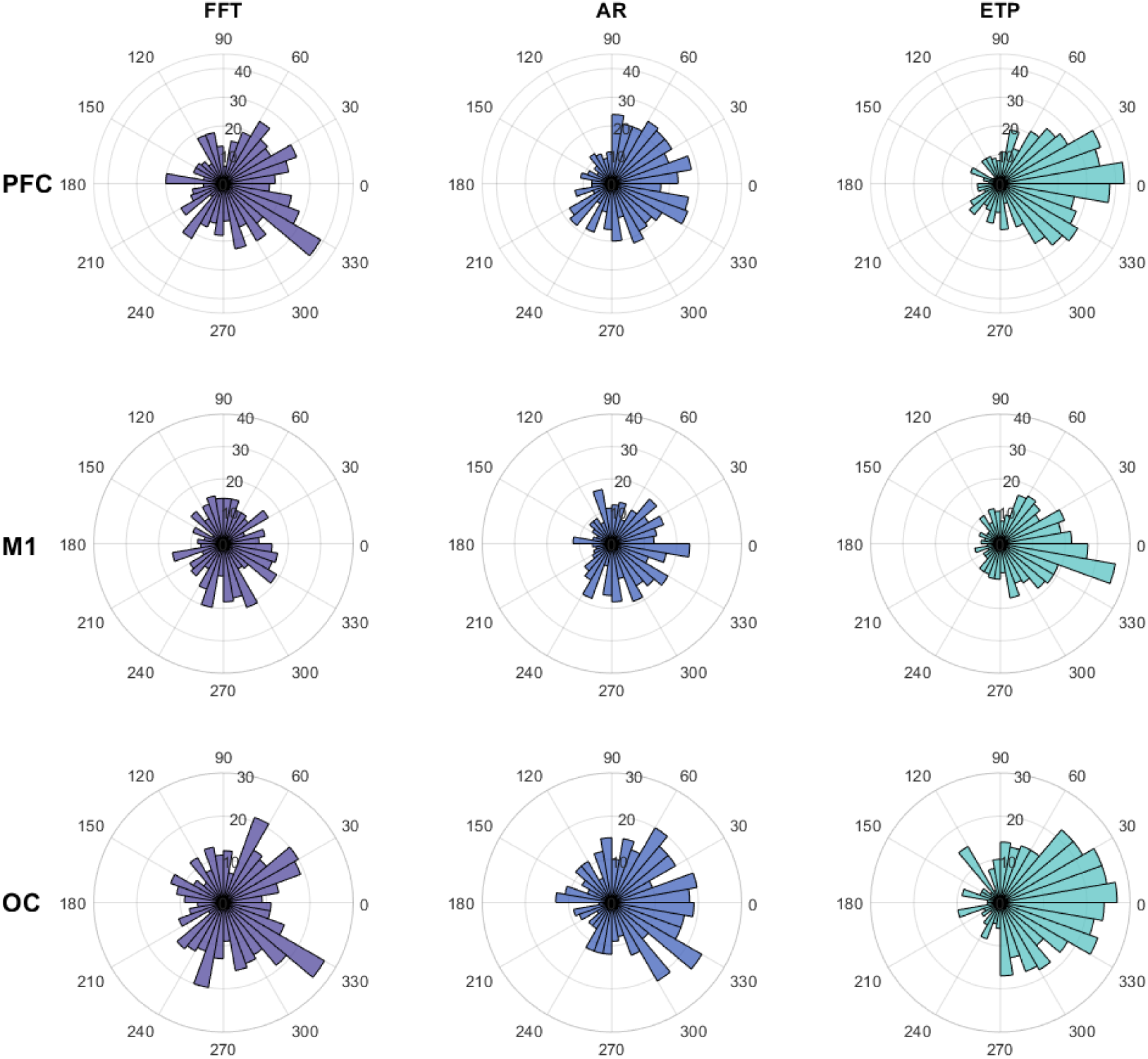
Phase estimation results for the real-time experiment for each algorithm and brain region in the beta band.

**Figure S8.**
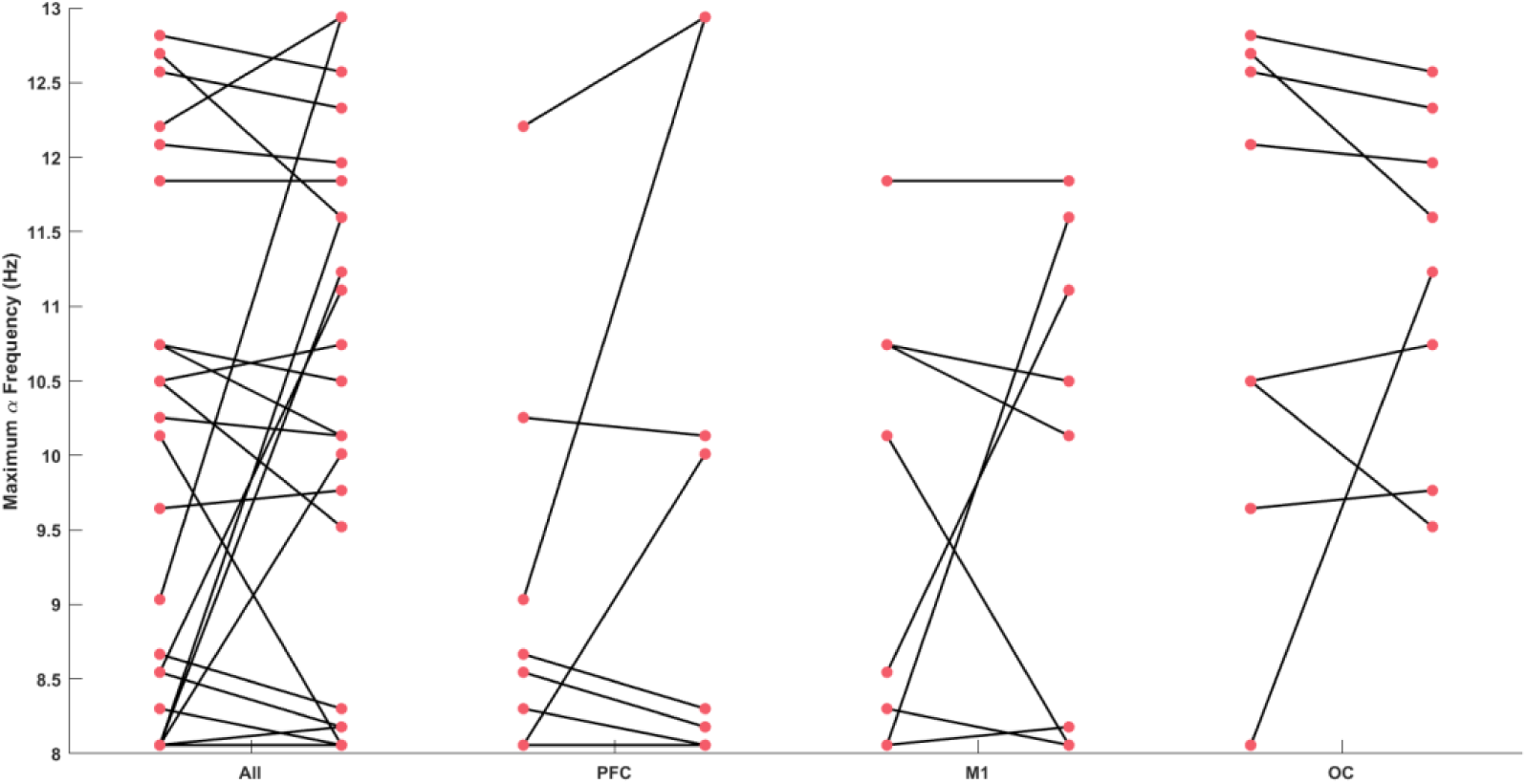
Individual’s maximum alpha frequency between pre- and post-experiment resting-state EEG over different regions.

